# Zebra finches use brightness cues for gap selection in flight

**DOI:** 10.1101/2023.02.21.529441

**Authors:** Emma Borsier, Graham K. Taylor

## Abstract

Flying animals that are adapted to life in cluttered environments require effective and robust guidance mechanisms to avoid collisions. Birds and insects use optic flow cues to avoid obstacles when flying, but these are only generated by self-motion and are likely to be used in conjunction with other cues such as visual contrast between foreground and background objects. Orchid bees use a strategy based on relative brightness to find clear paths through dense environments. To investigate the possibility that birds use a similar strategy, we presented zebra finches *Taeniopygia guttata* with symmetric or asymmetric brightness distributions behind the aperture of a tunnel and recorded their flight through it. The background brightness conditions influenced both the birds’ decision to enter the tunnel and their flight direction upon exit. Zebra finches were more likely to initiate flight through the tunnel if they could see a bright background through its aperture and were more likely to fly to the bright side on exiting the tunnel. We found no evidence of a centring response during gap negotiation; instead, the zebra finches entered the tunnel by turning tightly around its near edge. These results hint at a possible pre-planning of the trajectories before the onset of flight.

## Introduction

Flying animals achieve fine scale trajectory control to avoid collisions in cluttered environments. Birds and insects rely on vision to guide their flight, with many convergent similarities in the detailed mechanisms that they use. Many of the best-known visual guidance strategies of birds and insects are based on the use of optic flow (1–10), which describes the apparent motion of objects in the visual field resulting from the movement of an observer or its environment. One such behaviour, first described in honeybees *Apis mellifera* (5) and later reported in budgerigars *Melopsittacus undulatus* (6), involves steering flight to balance the optic flow between the left and right visual hemispheres. This response produces centred flight along a corridor with symmetrically striped walls, but it is far from clear what its effect would be in cluttered natural environments such as forests, or what other responses might be involved in negotiating such environments. Indeed, subsequent research has found no evidence for an optic-flow balancing response in hummingbirds *Calypte anna*, whose lateral steering behaviour instead depends on the vertical extent of visual features in the environment (11).

Natural environments such as forests can be densely cluttered, which makes the optic flow fields that they produce complex to interpret. Nevertheless, birds and insects manage to find their way quickly through clutter, so the guidance strategies they use must be robust to this complexity. One possibility is that they achieve this robustness by integrating other types of visual information with optic flow cues. For example, Baird and Dacke (12) found that orchid bees *Euglossa imperialis* employ a strategy based on using relative brightness cues to detect and negotiate gaps in their environment. When presented with apertures of different shapes or varying background lighting conditions, the orchid bees flew towards the point of greatest brightness within the aperture. It is not yet known whether birds employ a similar strategy to detect and negotiate gaps during flight, but it seems likely that they do, given the tendency of birds to fly towards windows when trapped in a room, and the tendency of nocturnal migratory birds to be disoriented by artificial sources of light (13–15).

Like orchid bees, many birds inhabit cluttered environments such as forests, in which brightness is a reliable indicator of the open space between darker obstacles. Interestingly, the visual system that guides landing in budgerigars relies on edge contrast (16), but it remains to be determined whether they use edge contrast only for landing, or more broadly for finding a path through clutter. Here we ask whether zebra finches *Taeniopygia guttata* use brightness cues for gap selection and negotiation in flight. To answer this question, we recorded the flight behaviour of captive birds when presented with varying background brightness conditions through the aperture of a narrow flight tunnel between two adjoining aviaries (Movie S1). Our results show that whether and how the birds fly through the tunnel is strongly influenced by the background lighting conditions, shedding new light on how birds use brightness cues to structure their flight behaviour.

## Materials and methods

### Animals

We used a captive population of *N*=24 adult zebra finches, comprising 13 males and 11 females. Birds could be sexed using characteristics visible in the video we collected but could not be individually identified.

### Experimental design

We conducted the experiments in a large aviary with multiple perches, comprising an interior and an exterior part connected by a square-section tunnel of width 0.33 m and length 0.58 m, through which the birds could fly freely (Fig. 1A; Movie S1). The interior aviary had white-painted walls and bright overhead LED lighting; the exterior aviary received natural lighting through its mesh frontage (see Supplementary Methods). We filmed *n*=419 flight trajectories of zebra finches travelling from the interior to the exterior through the tunnel, using two synchronised 4k cameras recording at 120 frames per second (E2, Z-CAM, Shenzhen, China). The cameras were mounted below or behind the entrance to the tunnel (Fig. 1A), aligned to the tunnel’s centreline. We recorded the birds’ flight behaviour over 12 days, recording two 1 to 1.5 h sessions per day. To test how destination brightness affected the birds’ flight trajectories, we attached two panels of black or white fabric side-by-side on the mesh, giving four experimental conditions: dark-dark, bright-bright, bright-dark and dark-bright (Fig. S1). We presented the symmetric dark or bright conditions on 5 of the 12 test days, alternating presentation of the asymmetric bright-left and bright-right conditions between sessions on the other 7 test days (Table S1).

**Figure 1.**
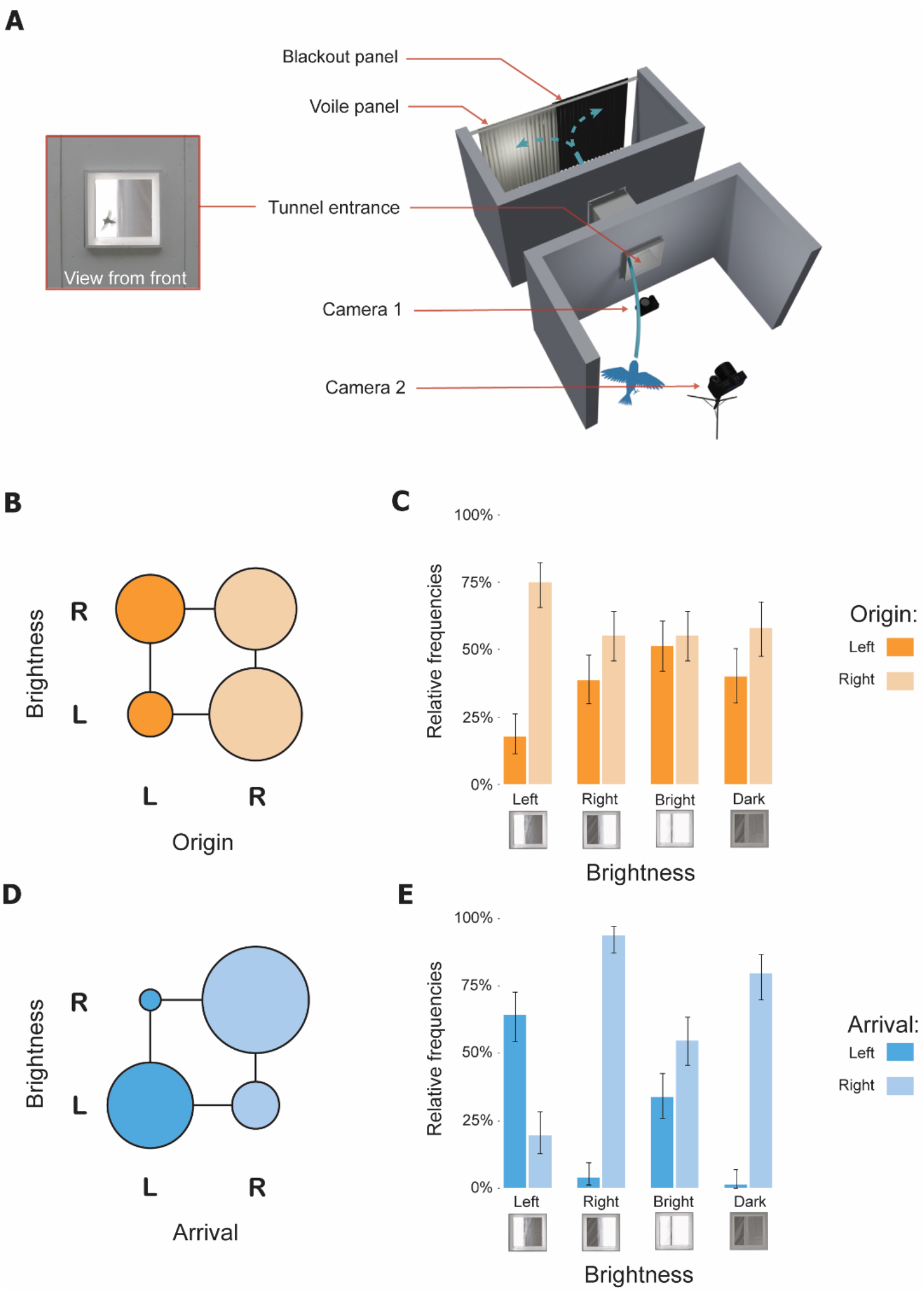
Flight of zebra finches through a tunnel between aviaries was influenced by the background brightness distribution visible through its aperture. **(A)** Representation of the experimental setup (not to scale), with Camera 2’s view of the tunnel (on the left, see Movie S1). **(B,D)** Bubble plots showing the number of flights that: **(B)** originated from; or **(D)** arrived on, the left (“L”) or right (“R”) side of the aviary, depending on whether the background was bright to the right (“R”) or bright to the left (“L”). Circle area is shown proportional to count. **(C,E)** Bar charts showing the frequency of: **(C)** origin from; or **(E)** arrival on, the left (“L”) and right (“R”) sides of the aviary, according to background brightness condition. The icons below the plots illustrate the appearance of the background behind the aperture, and frequencies shown are relative to the total number of flights recorded under each brightness condition (i.e. including those where the side of origin or arrival was ambiguous).

### Video analysis

For each flight, we recorded whether the trajectory originated from the left or right side of the interior aviary, and whether it arrived on the left or right side of the exterior aviary after exiting the tunnel (Table 1). We quantified the position of the bird’s entry point *φ* using the video collected from the camera placed below the tunnel entrance. We identified the frame on which the bill tip first became obscured by the lower lip of the tunnel. We then used FIJI v1.53C software to measure the radial projection of the bird’s bill onto the lower lip of the tunnel, normalised such that *φ* = −1 represents the left edge, *φ* = 0 the centre, and *φ* = 1 the right edge (Fig. S3). This quantity does not account for variation in the height at which the bird entered the tunnel, so is not linear in the bird’s lateral position. Nonetheless, because the camera was aligned to the centreline of the tunnel, negative entry point values always correspond to entry on the left whereas positive entry point values always correspond to entry on the right.

**Table 1.**
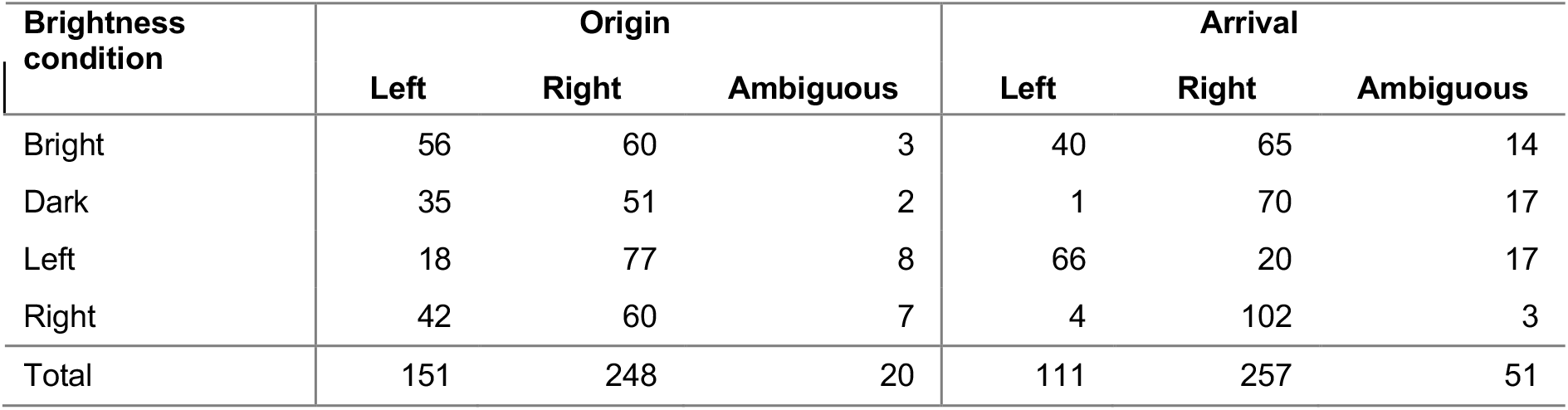
Frequency distributions of the side of Origin and of Arrival variables of the flight trajectories for each of the four different brightness conditions. The side of origin or arrival was sometimes ambiguous in flights with S-shaped trajectories.

| Brightness condition | Origin |  |  | Arrival |  |  |
| --- | --- | --- | --- | --- | --- | --- |
|  | Left | Right | Ambiguous | Left | Right | Ambiguous |
| Bright | 56 | 60 | 3 | 40 | 65 | 14 |
| Dark | 35 | 51 | 2 | 1 | 70 | 17 |
| Left | 18 | 77 | 8 | 66 | 20 | 17 |
| Right | 42 | 60 | 7 | 4 | 102 | 3 |
| Total | 151 | 248 | 20 | 111 | 257 | 51 |

### Statistical analysis

We analysed the data statistically using R (v4.0.3) and R Studio (v1.3.1093) software. As it was not possible to identify individuals from the video footage, we were limited to treating flight trajectories recorded from the same birds as independent data points. We report 95% Wilson score confidence intervals (CIs) for sample proportions and odds ratios, noting that these CIs are expected to have less than nominal coverage probability because of the nonindependence of flights by the same individuals. We used a two-tailed Mann-Whitney U test to assess whether the distribution of the birds’ entry positions varied between the two asymmetric brightness conditions, and used chi-square tests to assess whether the frequencies of origin and arrival on the left versus right sides of the aviary varied according to whether the bright side was on the left or the right.

## Results

We sampled a total of *N*=419 flight trajectories from a population of *n*=24 zebra finches (Table 1). Although the population sex ratio was approximately balanced (54% males), there was a strong bias towards males in the flights we recorded (88% males), with males approximately six times more likely to fly through the tunnel than females (odds ratio: 6.2; CI: 2.7, 14.5).

Most of the flight trajectories arrived on the right (70%; CI: 65, 74%; n=368 unambiguous records), presumably reflecting a preference for perches located on the right side of the exterior aviary (Table 1; Fig. 1D). This overall preference was nevertheless associated with a highly significant relationship between side of arrival and brightness condition (χ^2^(3) = 154.63, p < .001), with the birds switching their arrival side preference from right to left when the background was bright to the left (Table 1; Fig. 1E). Considering the two asymmetric brightness conditions together, the birds were seven times more likely to arrive on the bright side than the dark side (odds: 9.0; CI: 4.6, 10.7). It follows that the birds selected the goal of their flight using brightness cues.

Most of the flight trajectories originated from the right (62%; CI: 57, 67%; *n*=399 unambiguous records), which may again reflect a preference for perches located on the right side of the aviary (Fig. 1B). This bias was displayed under all four test conditions (Table 1; Fig. 1C), but there was still a statistically significant relationship between side of origin and brightness condition (χ^2^(3) = 20.567, *p* < .001). Specifically, the birds’ tendency to originate from the right was amplified when the background was bright to the left rather than bright to the right (odds ratio: 3.0; CI: 1.6, 5.7; Fig. 1C). Under this test condition, the tunnel aperture would have appeared bright when viewed from the right and dark when viewed from the left, consistent with the hypothesis that the birds were preferentially targeting brighter gaps.

The birds’ tunnel entry points did not appear to be centred under any of the test conditions: instead, the birds tended to enter the tunnel on either the left or the right, and only rarely in the middle (Fig. 2A). A Hartigan’s dip test confirmed that the entry point distribution was non-unimodal under each of the symmetric test conditions (bright: *D* = 0.072, *p* < .001; dark: *D* = 0.066, *p* < .01), and under the asymmetric test condition with the background bright to the right (*D* = 0.057, *p* < .01), where the first two estimated modes were at *φ* = 0.33 and *φ* = 0.35. For the asymmetric condition with the background bright to the left, there was no evidence for non-unimodality (*D* = 0.025, *p* = 0.950), with the mode estimated at an entry point of *φ* =0.27 (Table 2).

**Figure 2.**
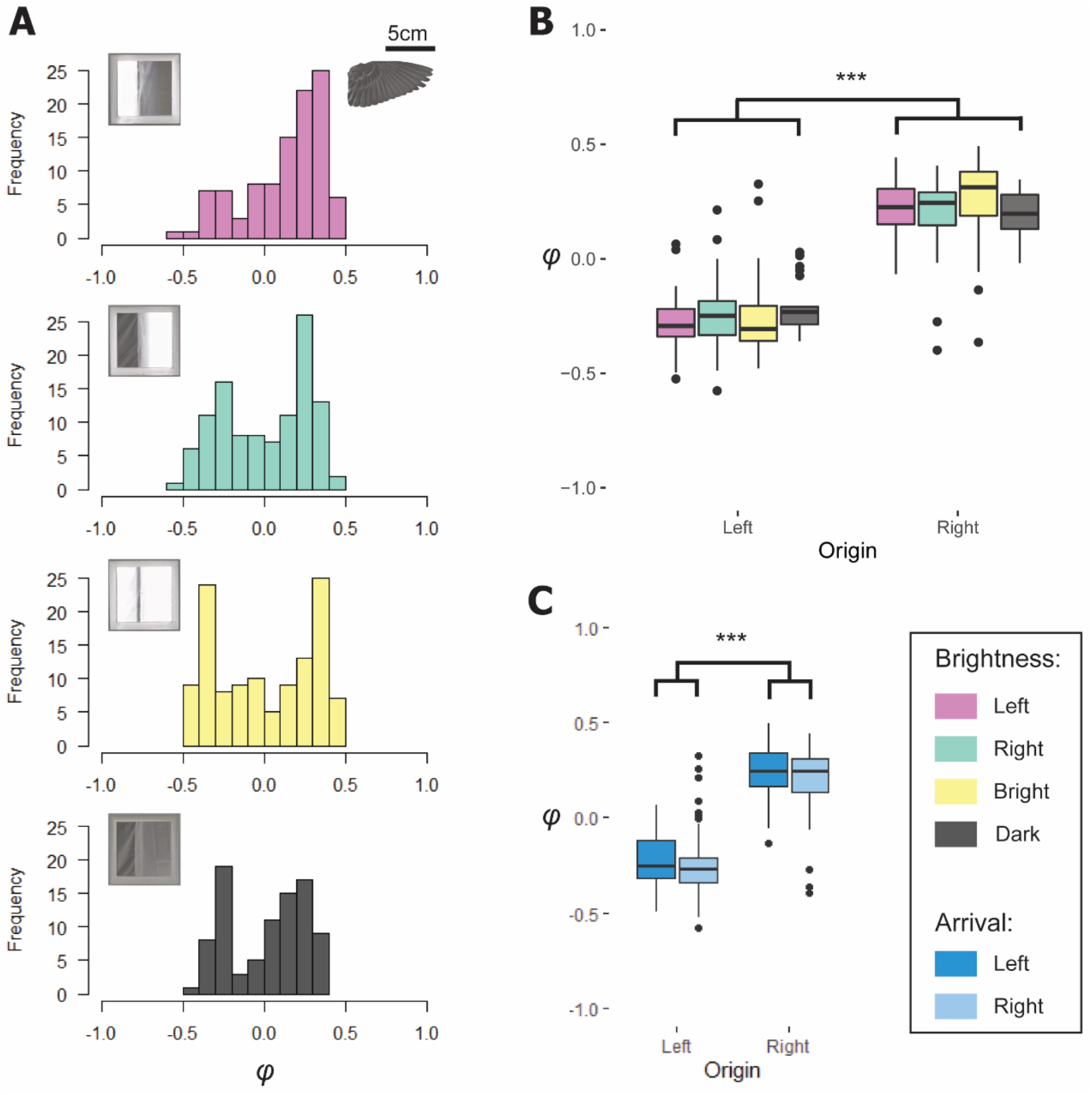
The position of the entry point of zebra finches into the tunnel depends only on the side they arrive from. **(A)** Histograms of entry-point position *φ* under each of the four background brightness conditions, as illustrated by the icons (from top to bottom: bright to the left, bright to the right, bright symmetric, dark symmetric). A zebra finch wing is depicted for scale, showing the approximate extent of the clearance. **(B,C)** Boxplots of entry-point position *φ* by side of origin, grouped according to: **(B)** background brightness condition; and **(C)** side of arrival.

**Table 2.** Modal entry point of the zebra finches as they entered the tunnel under each of the four brightness conditions.

| Brightness condition | Entry point mode(s) |
| --- | --- |
| Bright | -0.329, 0.353 |
| Dark | -0.246, 0.282 |
| Left | 0.266 |
| Right | -0.241, 0.267 |

The position of the birds’ point of entry into the tunnel was dependent on their side of origin (two-sample *t*-test: *t*(397) = −30.8, *p* < .001): birds originating from the left tended to enter to the left of centre (*φ* = −0.24 ± 0.15; mean ± s.d; *n* = 151), whereas birds originating from the right tended to enter to the right of centre (*φ* = 0.23 ± 0.15; mean ± s.d; *n* = 248). However, whereas the birds’ side of origin was itself related to brightness condition (see above), brightness condition had no statistically significant effect on the birds’ entry point over and above that already contained in the effect of origin, when comparing the two asymmetric brightness conditions (two-way ANOVA: *F*(1, 173) = 0.722, *p* = .397; Fig. 2B). In contrast, the side on which the birds arrived had a significant effect on the position of their point of entry into the tunnel when controlling for their side of origin (two-way ANOVA: *F*(1,345) = 4.98, *p* = .026; Fig. 2C). The birds’ point of entry into the tunnel was therefore determined by where they were flying from, and by where they were flying to, each of which in turn relates to brightness.

## Discussion

The results of this simple flight experiment (Fig. 1A) demonstrate that the flight trajectories of zebra finches are influenced by varying brightness conditions. Specifically, the distribution of background brightness seen through an aperture affects: (i) how likely a bird is to take off and fly through the aperture (Fig. 1 B,C); (ii) from which side the bird enters the aperture (Fig. 2A); and (iii) where the bird flies to after exiting (Fig. 1 D,E). Our results therefore confirm that birds use visual contrast to select their flight trajectories during tasks including take-off, gap negotiation, and perching.

Our zebra finches did not aim for the centre of the tunnel aperture upon entry (Fig. 2A). Instead, they entered the tunnel on the side they were perched before take-off (Fig. 2B,C), resulting in a bimodal entry-point distribution (Fig. 2A). Hence, our observations do not support the hypothesis that zebra finches centre their flight by balancing the optic flow between their left and right visual hemispheres, as has been observed in bumblebees (4), honeybees (5), and budgerigars (6). Instead, our zebra finches turned tightly around the near corner of the tunnel aperture, aiming at an entry point approximately 0.12 m from the near wall. This behaviour left them with a clearance of just over one wing length (Fig. 2A), consistent with results from Harris’ hawks, which also aim for a point giving a clearance of just over one wing length when steering around obstacles (17).

Altering the visual contrast that the birds could see through the tunnel aperture affected the side that the birds entered it from (Fig. 1C). Although the birds preferred to enter the tunnel from the right, they were three times more likely to do so if the background viewed through its aperture was brighter to the left than to the right (Fig. 1B). Since the lefthand side of the background was visible from the righthand side of the aviary, and vice versa, it follows that the birds were more likely to take off and fly through the tunnel if the tunnel appeared as a bright aperture from their original perching point. Likewise, when exiting, the birds were more likely to fly towards the brighter side of the outdoor aviary (Fig. 1D). This suggests that the goal of the birds’ flight behaviour was decided before take-off and was structured in relation to brightness cues.

Our findings from zebra finches are therefore broadly consistent with the findings from orchid bees by Baird and Dacke (12), who posited that in cluttered environments such as forests, brightness cues “provide information about the point of greatest clearance in an aperture, the relative size of an aperture as well as the clearest path” (12). However, whereas orchid bees appear to steer their flight using background brightness cues, targeting the brightest point of an unevenly lit aperture, we found no evidence that the lateral position of the bird’s entry point was affected in detail by the background brightness distribution (Fig. 2A), except insofar as this brightness distribution determined the side of the aviary from which they entered the tunnel (Fig. 2B).

Finally, it is worth noting that males were over-represented in our flight data. Because we could not identify individuals reliably between flights, the exact number of biological replicates is unknown, and we were unable to account for between-individual variation in our analysis. Budgerigars are known to adopt idiosyncratic flight trajectories for repetitive flight tasks (18), so controlling for between-individual variation in the flight trajectories could have provided further information on the effect of contrast for this experiment. Nonetheless, our results clearly show that brightness cues are important in determining how zebra finches choose to structure their flight when negotiating gaps in closed environments.

## Acknowledgements

We thank Helen Sanders and Lucy Larkman for husbandry. This project has received funding from the European Research Council (ERC) under the European Union’s Horizon 2020 research and innovation programme (Grant Agreement No. 682501). EB’s work was supported by funding from the Biotechnology and Biological Sciences Research Council (BBSRC) [grant number BB/T008784/1], via the Interdisciplinary Bioscience Doctoral Training Partnership.

## Ethics statement

This work received approval from the Animal Welfare and Ethical Review Board of the Department of Zoology, University of Oxford, in accordance with University policy on the use of protected animals for scientific research, permit no. APA/1/5/ZOO/NASPA, and is considered not to pose any significant risk of causing pain, suffering, damage or lasting harm to the animals. No adverse effects were noted during the trials.

## Notes

### Competing Interest Statement

The authors have declared no competing interest.

