## Supplementary Material for "Zebra finches use brightness cues for gap selection in flight"

### Supplementary Methods

#### Aviary

The dimensions of the cuboidal tunnel separating the indoor and outdoor aviaries were 58 cm in length, 33 cm in width, and 33 cm in height. The dimensions of the indoor and outdoor aviaries were both 5 m in length, 2 m in width, and 2.7 m in height (maximum dimensions). Feeders were set up in the outdoor aviary while nest boxes and a water dispenser were set up indoors to encourage the birds to fly through the tunnel regularly. Perches and nest boxes were evenly distributed on each side of the aviaries to minimise the preference of the birds for a given side. The indoor aviary was lit with a mixture of 6500 K and 4000 K flicker-free LED lighting, and the outdoor aviary received natural light through the metal mesh of the walls facing out (in front and to the right of the tunnel).

#### Experimental set-up

In the outdoor aviary, a pair of semi-transparent white voile panels were set up opposite from the tunnel, such that it is the only visible background when looking through the tunnel from all angles (see Figure 1A). This allowed the natural light to stream through the outdoor aviary and the tunnel, while removing any visual landmarks or potential asymmetry in the optic flow created by the left and right sides of the tunnel's exit. To block out the natural light on either or both sides of the tunnel in the outdoor aviary, we used blackout curtains which were set up behind the voile panels and attached to the metal mesh. As viewed from inside, the blackout curtains could appear to be either brighter or darker than the white wall of the interior aviary depending on weather conditions; the intensity of the natural light measured outside the tunnel varied from 500 to 8000 lux during the experiment. The brightness gradient of the background was never completely uniform due to the Sun’s position in the sky. During the experiment, the angle between the axis of the tunnel and the solar azimuth varied from -23 degrees (at the start of recording) to 40 degrees (at the end of recording).

#### Video recording

Two high-speed cameras (Z Cam E2 Cinematic Camera) were used for video recording. The cameras were recording at 120 frames per second with 3840 x 2160 pixel resolution, ISO 8000, 1/4000 shutter speed, and F/1.8 aperture. One camera was set up in the recessed wall opposite the tunnel’s entrance such that it was facing the tunnel and looking directly through it from a 220 cm distance, while having a wide view of the indoor aviary. The second camera was set up 110 cm below the entrance of the tunnel facing upwards, such that it had the entrance within frame as well as a view of the upper half of the indoor aviary. The recordings from both cameras were synchronised using the built-in multi-camera synchronisation feature.

The voile panels and blackout curtains were set up in the outdoor aviary 15 to 20 minutes before the first recording session every day, and were taken out after the last session. One brightness condition was used per recording session, and each of the four conditions were used for multiple sessions on separate days. The “Bright left” and “Bright right” asymmetric sessions were mutually paired and alternated each day such that one condition was always followed or preceded by the other. This was done to eliminate the possibility that any difference in the birds' behaviour between the sessions with asymmetric conditions could be attributed to behavioural changes specific to the day, time of day or the order of the sessions.

### Supplementary figures and tables

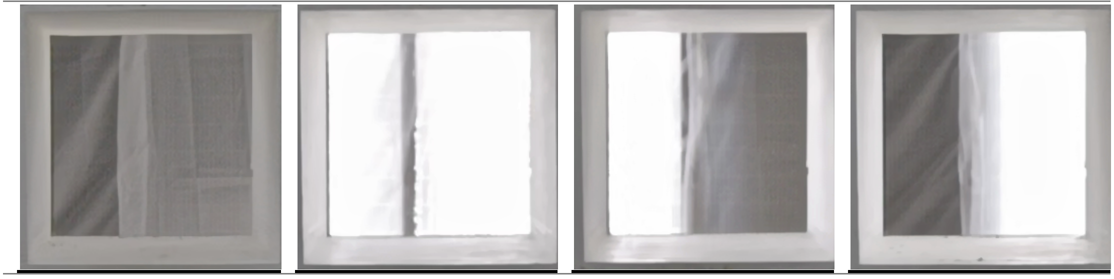


**Figure S1.** Photos of the tunnel and the view of the background for each of the four brightness conditions: A) Dark background; B) Bright background; C) Brighter on the left; D) Brighter on the right.

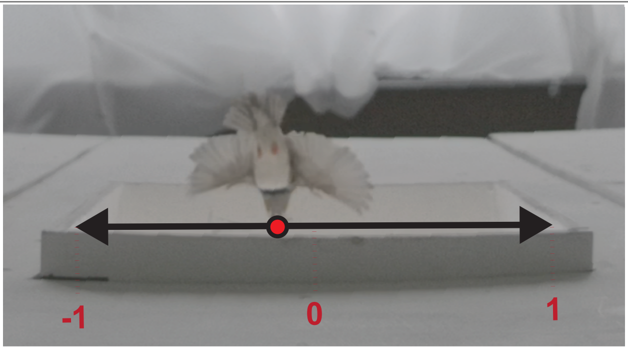


**Figure S2.** Example of an “entry” frame with a representation of the variable measured, i.e. the horizontal position of the bird, measured as it enters the tunnel, at the tip of the beak (highlighted in red).

**Table S1**. Distribution of the flight data across the four contrast configurations. The number of sessions, number of days and hours of footage covered in the analysis are shown for each brightness condition.

| Brightness condition | Number of sessions | Number of days | Hours of footage | Number of individual flights |
|---|---|---|---|---|
| Bright | 4 | 2 | 2.5 | 119 |
| Dark | 6 | 3 | 3.6 | 88 |
| Brighter on right | 7 | 7 | 7.1 | 103 |
| Brighter on left | 7 | 7 | 7.7 | 109 |
| All combined | 24 | 12 | 21.0 | 419 |
